# Circadian feeding rhythm loss drives sex-specific β-cell dysfunction in diet-induced obesity

**DOI:** 10.64898/2026.11.02.763239

**Authors:** Luhui Zhang, Satish K. Sen, Kuntol Rakshit, Thanh Nguyen, Kazuno Omori, Ananya Bharath, Zong Wei, Alexandre Gaspar-Maia, Aleksey V. Matveyenko

## Abstract

Diet-induced obesity (DIO), modeled by a high-fat diet (HFD), adversely affects circadian glucose homeostasis partly by dysregulating fasting/feeding cycles. However, its impact on circadian β-cell function and relevant transcriptional programs remains unclear. To investigate this, we exposed male and female C57BL/6 mice to either chow or HFD for eight weeks and assessed behavioral (feeding/activity), physiological (glucose tolerance/insulin secretion), transcriptomic (RNA sequencing (RNA-seq)), and epigenomic (Assay for Transposase-Accessible Chromatin using Sequencing (ATAC-seq)) circadian rhythms. Additionally, we tested whether normalizing fasting/feeding cycles via time-restricted feeding (tRF) during the dark- or light-phase of the circadian cycle could mitigate DIO’s deleterious effects. DIO disrupted circadian fasting/feeding rhythms, with greater disruption in males than females (p<0.05 vs. CHOW). In males, DIO abolished circadian regulation of glucose tolerance and *in vivo* glucose-stimulated insulin secretion (GSIS) (p<0.05 vs. chow), whereas female mice maintained circadian glucose homeostasis. RNA-seq analysis of male islets revealed a 42.1% reduction in circadian-regulated transcripts under DIO, with weakened rhythms in key β-cell pathways such as insulin secretion. Moreover, ATAC-Seq analysis indicated reduced chromatin accessibility of genes/pathways related to insulin secretion and glucose hemostasis under DIO (p<0.05 vs. chow) during both dark- or light-phase of the circadian cycle. Notably, dark-phase time-restricted feeding (tRF) restored circadian glucose tolerance, GSIS, and islet gene expression, whereas light-phase tRF enhanced β-cell function during the normally inactive/light phase of the circadian cycle, further suppressing circadian rhythmicity of the islet transcriptome. Motif analysis indicated that DIO suppressed the activity of D-box transcription factors (TFs) (DBP and TEF) while enhancing the activity of ETS family TFs (ELK1, ETS2, ELF2, and GABPA), which was partly mitigated by dark-phase tRF. Our study demonstrates that DIO disrupts circadian β-cell function and gene expression by dysregulating fasting/feeding cycles in male mice and reveals significant sex differences in how DIO affects circadian glucose homeostasis and β-cell function.

## Introduction

Obesity and type 2 diabetes (T2DM), characterized by insulin resistance and β-cell dysfunction, are among the most prevalent metabolic disorders worldwide, and their incidence continues to rise in parallel(*1, 2*). Insulin resistance alone, however, is not sufficient to produce hyperglycemia: the great majority of insulin-resistant individuals maintain normoglycemia through compensatory increases in insulin output, and type 2 diabetes emerges when pancreatic β cells fail to sustain that compensation(*3, 4*). β-cell dysfunction is therefore the proximate lesion in the transition from obesity to type 2 diabetes and is characterized by impaired glucose-stimulated insulin secretion, loss of first-phase secretory kinetics, and disruption of the transcriptional programs that maintain β-cell identity(*5, 6*). Understanding how chronic nutrient excess compromises β-cell function is consequently central to understanding how obesity contributes to increased susceptibility for T2DM.

Critically, β-cell secretory function and corresponding glucose tolerance are not constant throughout the day and show strikingly robust time-of-day variations that favor enhanced insulin secretion and β-cell responsiveness at the onset of the active circadian phase(*7, 8*). This process has been shown be in part regulated by the core circadian clock, which exerts control over insulin exocytosis, glucose sensing, cell proliferation, response to various stressors, and the expression of core β-cell identity genes(*8–15*). On the molecular level, circadian clocks are cell-intrinsic transcriptional-translational feedback loops that align physiology with the 24-hour light-dark cycle(*16*). In the core loop, the bHLH-PAS activators CLOCK and BMAL1 drive transcription of *Per* and *Cry* genes through E-box elements, and the resulting PER-CRY complexes feedback to repress their own activators(*17*); an interlocking loop in which REV-ERB and ROR nuclear receptors compete at ROR response elements confers robustness of the circadian system and sets the phase for *Bmal1* expression(*18, 19*). Critically for transcriptional output in pancreatic β-cells, these core elements act combinatorially with a third class of element, the D-box, bound by the PAR-bZIP activators DBP, TEF, and HLF and antagonized by the repressor NFIL3(*10, 20*). E-box-, ROR-element-, and D-box-driven transcription peak at distinct times of day, and their combination is what distributes clock-controlled genes across the full 24-hour cycle rather than confining them to a single phase(*21*). Thus, metabolic tissues, including liver, adipose, skeletal muscle, and pancreatic islets, rely on this local circuitry to time nutrient handling to the fasting and feeding phases of the day(*11, 22–25*). Importantly, genetic disruption of the β-cell circadian clock in rodents or exposure to circadian disruption in humans is sufficient to impair insulin secretion, disrupt the adaption to obesity, and increased risk of T2DM(*13, 26, 27*).

There is a complex bidirectional relationship between obesity and the regulation of circadian rhythms. DIO has been shown to disrupt circadian behavior and gene expression in male mice, in part by disrupting the normal temporal restriction of feeding to the active circadian phase and thereby removing the principal entraining signal for peripheral clocks(*28, 29*). How this behavioral disruption manifests at the transcriptional and epigenetic level has been investigated most thoroughly in the mouse liver. DIO has been reported to dampen the rhythms of core clock genes (*Bmal1*, *Clock*, and *Per2*)(*28*), and/or also impair CLOCK:BMAL1 recruitment to canonical chromatin targets in mouse liver(*30*). Comparable effects of obesity on circadian organization are also evident in humans. Human obesity is also associated with disrupted feeding patterns and dampened clock gene expression in peripheral tissues(*31–35*). Importantly, diurnal rhythms in glucose tolerance and β-cell function appear to be either significantly attenuated or completely absent in obese individuals, as well as in patients diagnosed with T2DM(*12, 36*). However, the impact of obesity on circadian regulation of pancreatic islet function and genome-wide circadian transcriptional and chromatin-accessibility landscape remains unknown.

A further limitation of the existing literature is that the great majority of mechanistic studies of circadian metabolic dysfunction, including studies related to islet physiology, have been performed exclusively in males(*37*). Indeed, evidence supports distinct sex differences in the regulation of circadian rhythms with direct implications for metabolic health(*38, 39*). For instance, research findings reveal robust sex differences in the regulation of behavioral (sleep/activity cycles)(*40, 41*), hormonal (e.g., cortisol and melatonin secretion)(*42, 43*), metabolic (glucose tolerance)(*44, 45*), and molecular (tissue gene expression) circadian rhythms(*39*) underscoring the importance of assessing sex differences in circadian research. Furthermore, sex differences also permeate islet physiology with recent studies highlighting enhanced β-cell resilience to diabetogenic stress in female islets(*46*). Here, we combined behavioral, physiological, transcriptomic, and epigenomic profiling to identify novel mechanisms by which DIO disrupts circadian β-cell physiology and transcriptional/epigenetic control, and whether this effect is influenced by biological sex. We further used dark- and light-phase time-restricted feeding in HFD-fed male mice to determine whether directly correcting the diet-induced disruption of circadian feeding rhythm is sufficient to restore circadian β-cell physiology, and whether the circadian phase of the imposed feeding schedule determines the outcome of the intervention.

## Results

### High-fat diet disrupts circadian feeding behavior in a sex-specific manner

Circadian disruption of feeding behavior is a hallmark of DIO(*28*), but whether such disruption differs by sex, and how it relates downstream to islet physiology and gene regulation, remains poorly defined. To address this, we fed male and female C57BL/6 mice either standard chow or a high-fat diet (HFD) for 8 weeks and monitored feeding and locomotor activity by continuous recording during the last 2 weeks of the experimental period, followed by physiological testing and RNA-seq/ATAC-seq profiling of pancreatic islets (Fig. 1A). As expected, HFD feeding increased body weight in both sexes relative to chow-fed controls (p < 0.05), confirming that the DIO model was effective in both males and females (Fig. 1B, C). Constituently, HFD feeding also increased liver and epididymal white adipose tissue (eWAT) weight (p < 0.05), and elevated blood glucose and plasma insulin levels during both the light and dark phases of the circadian cycle (p < 0.05), in both males and females relative to chow-fed controls (Fig. S1A-J). To determine whether DIO altered the temporal organization of feeding behavior, we generated actograms of daily food intake. HFD blunted the light-dark cycle feeding rhythm in male mice, increasing the proportion of food intake consumed during the light (inactive) phase from 17.1% to 34.1% (*p*<0.05) at the expense of dark-phase intake (Fig. 1D-G). Interestingly, female mice on HFD largely preserved their nocturnal feeding pattern, with actograms, light/dark food intake distribution, and daily caloric intake resembling chow-fed females (Fig. 1H-K). In contrast to this disruption in feeding rhythm, HFD did not alter the overall pattern of locomotor activity rhythms in either sex (Fig. 1L-N, P-R), however female HFD-fed mice showed higher total daily activity than chow-fed females (p < 0.05) (Fig. 1O, S). Together, these behavioral data indicate that HFD selectively disrupts the circadian feeding rhythm in male mice without altering circadian locomotor activity in either sex, establishing a feeding-rhythm-specific, male-biased behavioral phenotype.

**Fig. 1.**
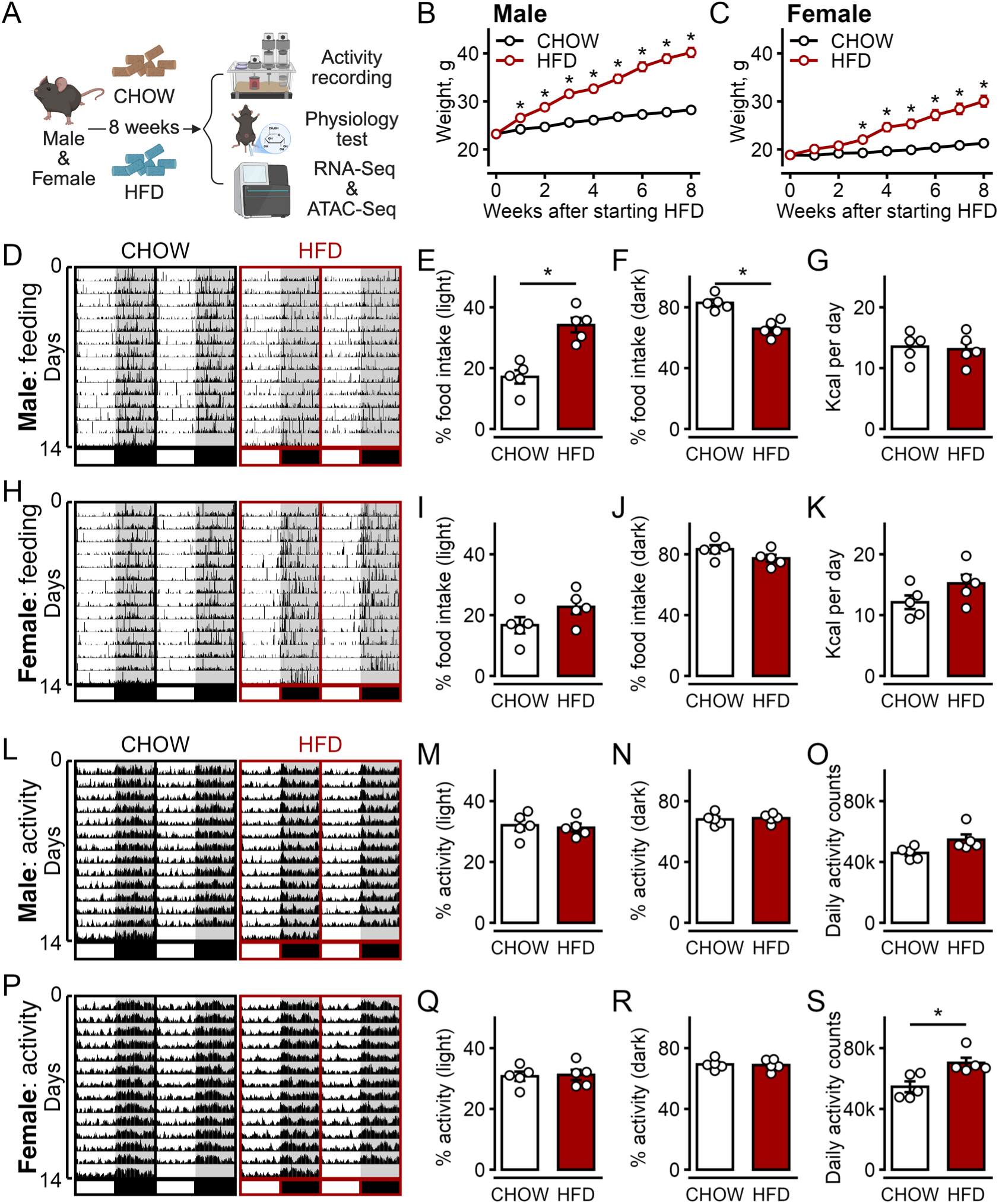
HFD feeding disrupts circadian feeding rhythm in sex-specific manner. (A) Schematic of experimental design. Male and female mice were fed chow or high-fat diet (HFD) for 8 weeks, followed by feeding and locomotor activity recording, metabolic phenotyping, and islet RNA-seq and ATAC-seq. (B, C) Body weight over 8 weeks in male (B) and female (C) mice on chow or HFD. n=18 mice per group. (D-G) Feeding behavior in male mice: double-plotted feedograms, averaged across 5 mice (D), proportion of daily food intake consumed during the light (E) and dark (F) phases, and total daily caloric intake (G). (H-K) Feeding behavior in female mice, as in D-G. (L-O) Locomotor activity in male mice: double-plotted actograms, averaged across 5 mice (L), proportion of daily activity during the light (M) and dark (N) phases, and total daily activity counts (O). (P-S) Locomotor activity in female mice, as in L-O. n=5 mice per group. White and black bars below feedograms/actograms indicate the light and dark phases of the circadian cycle, respectively. For (B) and (C), results were compared by two-way ANOVA with Fisher’s LSD test for post-hoc comparisons. For (E-G, I-K, M-O, Q-S), results were compared by unpaired two-tailed Student’s t test. Data are mean ± SEM; *p < 0.05.

### HFD impairs circadian glucose tolerance and glucose-stimulated insulin secretion in male mice

Because circadian feeding behavior gates the timing of nutrient exposure to peripheral tissues(*47*), we next tested whether the male-specific loss of feeding rhythmicity was accompanied by a parallel loss of circadian glucose homeostasis. We performed glucose tolerance tests (GTT) and assessed glucose-stimulated insulin secretion (GSIS) during the light and dark phases in each sex and diet group. In chow-fed male mice, glucose tolerance was enhanced in the dark phase compared to the light phase (p < 0.05), consistent with normal circadian gating of glucose handling(*10*). Importantly, diurnal difference in glucose tolerance was lost in HFD-fed male mice, such that glucose tolerance became uniformly impaired across the day and night compared to chow (p < 0.05) (Fig. 2A-D). In contrast, female mice maintained a light/dark difference in glucose tolerance on both chow and HFD (p < 0.05), indicating that circadian regulation of glucose tolerance was partially preserved despite DIO (Fig. 2E-H). *In vivo* GSIS mirrored this pattern: HFD abolished the light/dark difference in insulin secretion and its fold-change response in males, flattening what was a robust circadian rhythm in chow-fed males (Fig. 2I-L), whereas females retained circadian regulation of GSIS regardless of diet (p < 0.05) (Fig. 2M-P). Particularly, HFD decreased male GSIS primarily during the dark phase, reducing the averaged insulin fold-change response significantly from 2.29 in chow to 1.11 in HFD (p < 0.05) (Fig. 2L). As a measure of insulin action, we also performed insulin tolerance tests (ITT). Unlike glucose tolerance and GSIS, ITT did not show a light/dark difference in either chow- or HFD-fed male mice (Fig. S2A, B); instead, HFD decreased overall insulin sensitivity relative to chow in males (p < 0.05) (Fig. S2C, D), an effect that was not observed in females, whose insulin sensitivity was not affected significantly by HFD (Fig. S2E-H). Together, these results demonstrate that HFD disrupts both the circadian component of glucose homeostasis, as measured by GTT and GSIS, and overall insulin sensitivity, as measured by ITT, in males, whereas females are protected on both fronts, closely paralleling the sex-specific behavioral phenotype described above and raising the question of what molecular changes in the islet underlie this male-specific vulnerability.

**Fig. 2.**
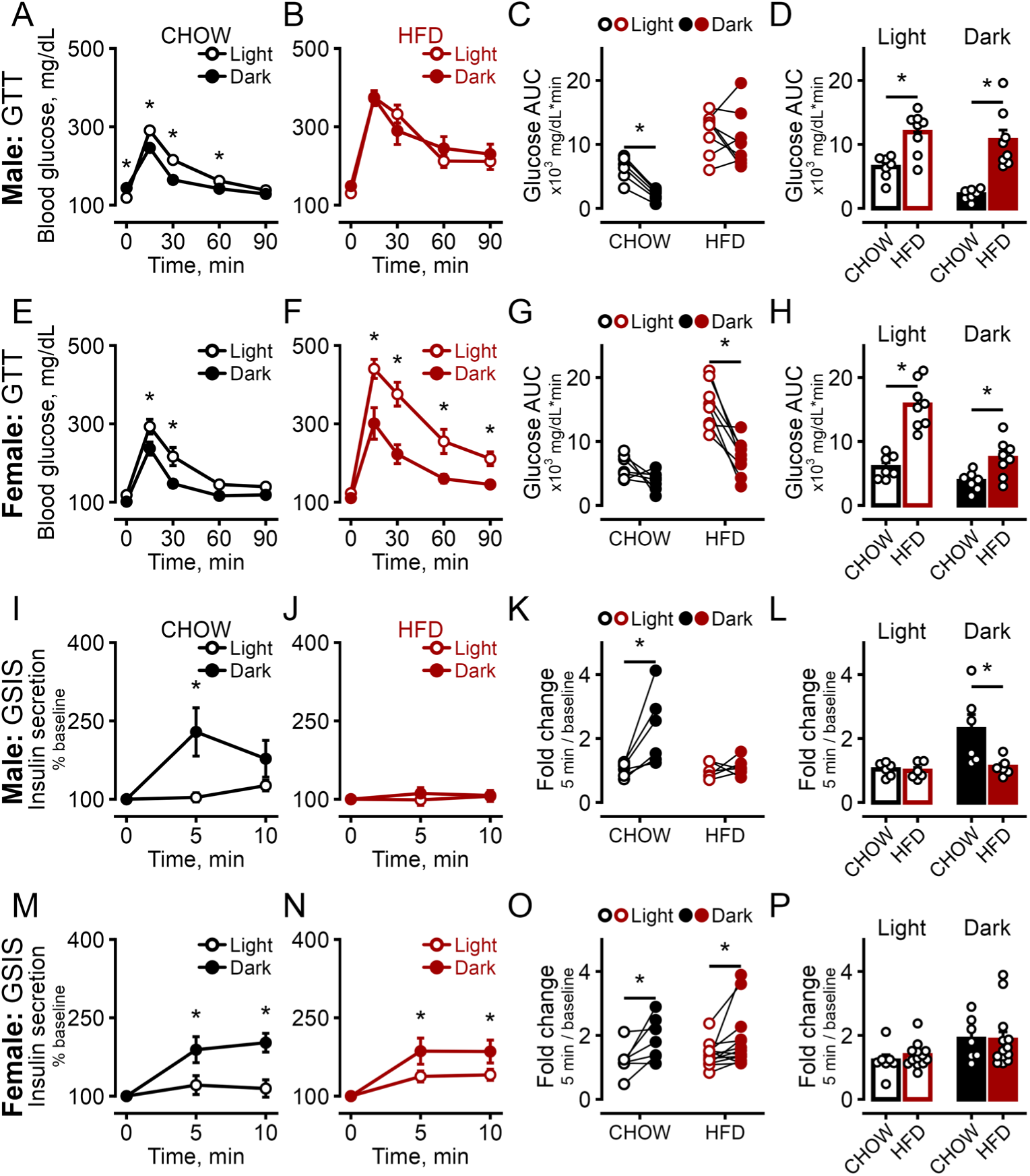
HFD feeding disrupts circadian rhythms in glucose tolerance and glucose-stimulated insulin secretion in sex-specific manner. (A-D) GTT in male mice. Blood glucose during an intraperitoneal GTT performed in the light or dark phase in chow-fed (A) and HFD-fed (B) mice; glucose area under the curve (AUC) by light/dark phase within each diet (C); and AUC compared between diets within each phase (D). n=7 mice for CHOW; n=8 mice for HFD. (E-H) GTT in female mice, as in A-D. n=7 mice for CHOW; n=8 mice for HFD. (I-L) GSIS in male mice. Plasma insulin (% of baseline) after intraperitoneal glucose administration in the light or dark phase in chow-fed (I) and HFD-fed (J) mice; fold change in insulin secretion (5 min relative to baseline) by light/dark phase within each diet (K); and fold change compared between diets within each phase (L). n=6 mice for CHOW; n=6 mice for HFD. (M-P) GSIS in female mice, as in I-L. n=7 mice for CHOW; n=13 mice for HFD. All results were compared by two-way ANOVA with Fisher’s LSD test for post-hoc comparisons. Data are mean ± SEM; *p < 0.05.

### HFD remodels the circadian islet transcriptome in male mice

To identify the molecular basis of this male-specific disruption, we performed circadian RNA-seq on islets from chow- and HFD-fed male mice collected across the 24-h cycle, focusing on genes with rhythmic expression. Circadian rhythmicity of gene expression was evaluated using the ARSER algorithm(*48, 49*). Direct comparison of the two conditions showed that 3,179 genes were rhythmic in chow-fed islets versus only 1,846 in HFD-fed islets, corresponding to a 42.1% overall reduction in circadian-regulated transcripts, with 579 genes shared between the two rhythmic gene sets (rhythmicity cutoff: FDR < 0.1) (Fig. 3A-C). Of the chow-rhythmic genes, 2600 lost rhythmicity under HFD (Fig. 3D); this cluster, exemplified by the representative gene *Adcy9* (Fig. 3E), was enriched for the Gene Ontology term related to key β-cell function, such as “insulin secretion” (Fig. 3F), and key genes within this pathway, including *Mafa*, *Hnf1a*, *Hnf4a*, and *Pck2*, also showed reduced amplitude under HFD, with modest phase shifts evident in a subset of these genes (Fig. 3G), indicating that HFD disrupts the circadian transcriptional program governing insulin-secretion machinery. A second set of 579 genes, including the key clock-controlled gene *Dbp* (Fig. 3H, I), remained rhythmic under both dietary conditions and was enriched for the GO terms including “circadian rhythm” (Fig. 3J), although the amplitude of 9 out of the 12 key circadian clock genes was reduced by HFD (Fig. 3K), suggesting that the male core circadian clock persists under DIO but with dampened rhythms. In addition, a third cluster of 1,263 genes gained rhythmicity under HFD that were not rhythmic in chow (Fig. 3L). This HFD-induced rhythmic cluster, exemplified by *Irak2* (Fig. 3M), was enriched for several stress-response pathways, including “positive regulation of NF-κB activity” (Fig. 3N), with core genes within this pathway showing increased amplitude under HFD besides becoming more rhythmic, such as *Irak1*, *Irak2*, *Npm1* (Fig. 3O), consistent with the emergence of a rhythmic islet stress-response program under DIO that is absent in the chow state.

**Fig. 3.**
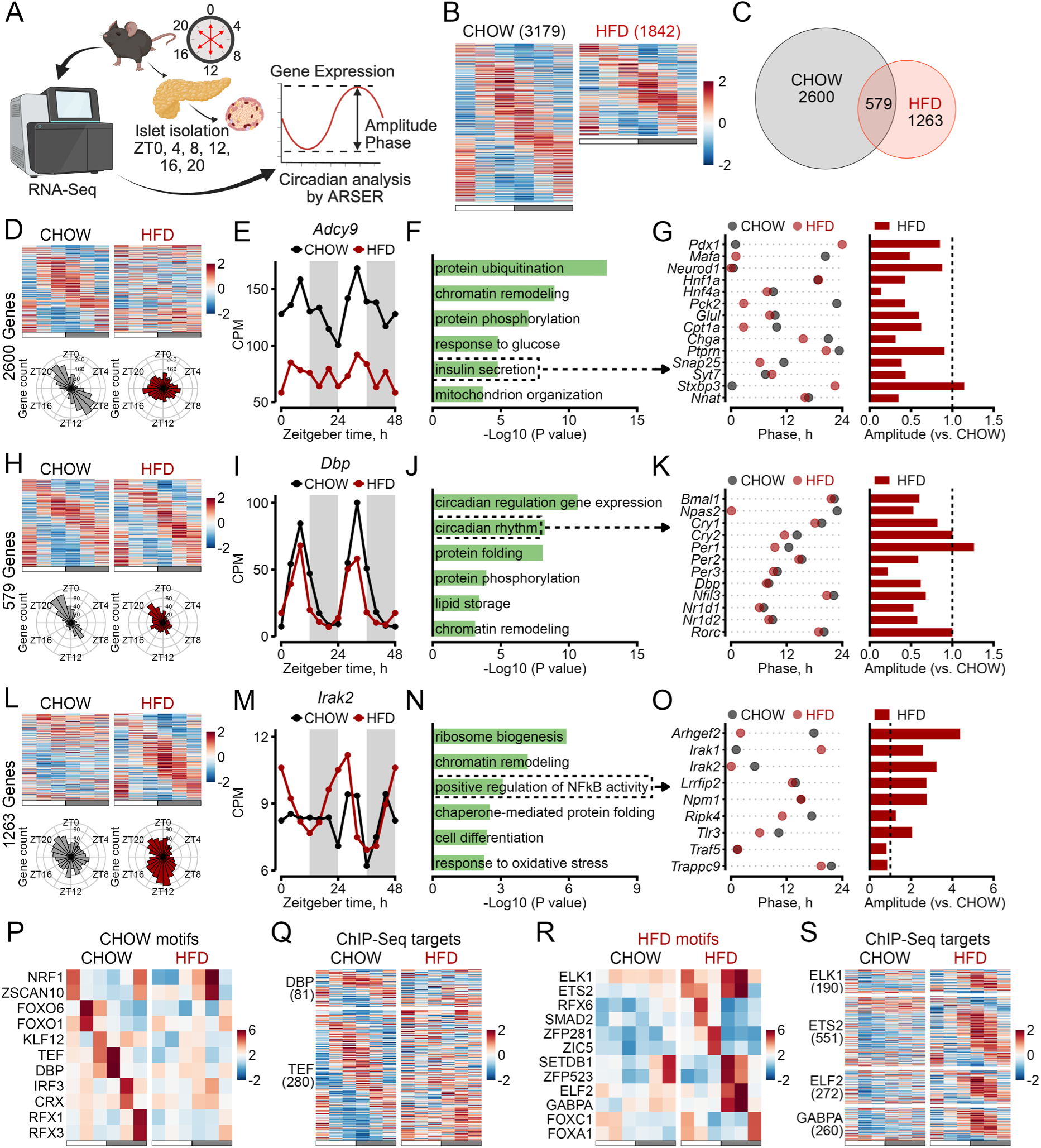
HFD feeding disrupts circadian rhythms in gene expressions of male mice. (A) Schematic of the study design. Islets were isolated for RNA-seq at 4-h intervals across the 24-h circadian cycle, with two independent replicates (mice) collected at each time point. Circadian rhythmicity of gene expression was assessed using ARSER with FDR cutoff at 0.1(*48, 49*). (B) Heatmaps of rhythmic genes in chow-fed (3,179 genes) and HFD-fed (1,846 genes) mice across the 24-h circadian cycle. Columns represent Zeitgeber time (ZT) points sampled every 4 h (from ZT0 to ZT20), with values averaged from two replicates. Gene expression was z-score normalized across CHOW and HFD groups. Genes are ranked by their peak-expression phase. White and gray bars below the heatmaps indicate the light and dark phases of the circadian cycle, respectively. (C) Venn diagram of the two gene sets, with 579 genes rhythmic under both diets. (D-G) Genes rhythmic in chow-fed but not HFD-fed mice (2,600 genes). Heatmaps of expression across the 24-h cycle with polar plots of peak-phase distribution in each diet (D); expression across Zeitgeber time for the representative gene *Adcy9* (E); Gene Ontology enrichment for the cluster, with “insulin secretion” highlighted (F); and peak phase and amplitude (G) of insulin-secretion genes in HFD relative to chow. (H-K) Genes rhythmic in both diets (579 genes), presented as in D-G, with the representative clock gene *Dbp* (I) and “circadian rhythm” highlighted (J, K). (L-O) Genes rhythmic in HFD-fed but not chow-fed mice (1,263 genes), presented as in D-G, with the representative gene *Irak2* (M) and “positive regulation of NF-κB activity” highlighted (N, O). For (E, I, and M), data from the two replicates per time point were plotted sequentially over a 48-h period, with replicate 1s of each time point displayed from 0-20 h and replicate 2s displayed from 24-44 h. The data point at 48 h is identical to that at 0 h and is included to facilitate visualization of circadian rhythmicity. (P, Q) Chow-enriched motifs identified via RcisTarget: transcription factor motifs enriched in circadian rhythmic genes in chow-fed mice (P); expression across the 24-h cycle of DBP and TEF ChIP-seq target genes that lose rhythmicity under HFD (Q). (R, S) HFD-enriched motifs identified via RcisTarget: motifs enriched in circadian rhythmic genes in HFD-fed mice (R); ELK1, ETS2, ELF2, and GABPA ChIP-seq target genes that gain rhythmicity under HFD (S). ChIP-seq targets of DBP were retrieved from Brown et al. (2021)(*10*), while targets of TEF, ELK1, ETS2, ELF2, and GABPA were obtained from the ChIP-Atlas database(*87*). Only target genes with binding sites within 1 kb of the transcription start site (TSS) were retained. For ChIP-Atlas datasets, targets were further required to exhibit a MACS2 binding score of at least 20 in ≥50% of samples in ChIP-Atlas. Color scales in (B, D, H, L, Q, and S) depict z-scored gene expression, whereas those in (P and R) depict normalized enrichment scores (NES) from RcisTarget.

To identify candidate transcriptional regulators underlying these changes, we performed motif enrichment analysis using RcisTarget(*50*) on the circadian rhythmic genes. Rhythmic genes were grouped into six clusters according to their peak expression phase, with cluster centers spaced at 4-hour intervals from ZT0 to ZT20, and motif enrichment analysis was performed separately for each cluster. Consistent with previous studies(*10*), we identified the D-box transcription factors DBP and TEF as highly enriched in chow-associated rhythmic genes with peak gene expression during the transition from the inactive to active circadian phase (e.g. ZT 8-12) (Fig. 3P). Importantly, enrichment for DBP/TEF motifs was greatly dampened in HFD-associated rhythmic genes highlighted by loss of rhythmic expression of previously identified DBP/TEF target genes (Fig. 3P, Q), specifically genes involved in the regulation of insulin secretion (*Snap25*, *Kcnh6*, *Slc30a8, Syt7, Stxbp3*), associated to obesity/diabetes (*Klf11*, *Kctd15*), insulin signaling (*Pik3r3*, *Ncoa5*), and glucose metabolism (*Pdk4*, *Gpd2*) (Fig. S3A). Moreover, exposure to HFD also led to the loss of motif enrichment for other key β-cell transcriptional regulators such as FOXO1(*51*) and RFX3(*52*). Conversely, the ETS-family transcription factors ELK1, ETS2, ELF2, and GABPA were enriched in HFD-associated regulatory regions, with their corresponding downstream target genes showing *de novo* rhythmicity under HFD in male islets (Fig. 3R, S). Notably, genes showing *de novo* rhythmicity under HFD included ER/mitochondrial stress genes (*Herpud1*, *Derl1*, *Eif2s1*, *Bag5*, *Dnaja3*) and inflammation-related genes (*Ptges3*, *Il17re*, *Sharpin*, *Cebpa*, *Dusp5*, *Irak1*) (Fig. S3B), implicating this transcription factor family in the HFD-induced stress-response rhythmic program. Together, these transcriptomic and motif enrichment data indicate that HFD both weakens the core circadian and insulin-secretion transcriptional programs and activates a new rhythmic stress-response program in male islets, potentially reflecting a shift in dominant transcriptional regulation from D-box factors toward ETS-family factors.

### HFD produces changes in diurnal islet chromatin accessibility indicating a heightened stress response

Because transcriptional rhythms are shaped by the underlying chromatin state, we next asked whether these diet-induced transcriptional changes were accompanied by altered chromatin accessibility. We performed ATAC-seq on male islets at ZT16 (dark phase) to capture chromatin accessibility changes at the time when β cells lost their GSIS ability significantly under HFD (Fig. 2L). At ZT16, 5,643 peaks were more accessible in chow relative to HFD (FDR < 0.1) (Fig. 4A), exemplified by *Bmal1* and *Dbp* (Fig. 4B); these chow-enriched regions were associated with genes in insulin-secretion-related pathways (Fig. 4C) and composed of genomic regions containing DBP (172 sites) and TEF (166 sites) motifs (Fig. 4D). Conversely, 4,513 peaks were more accessible in HFD relative to chow at ZT16 (FDR < 0.1) (Fig. 4E), exemplified by *Ralgapa2*, a negative regulator of glucose homeostasis whose deletion improves glucose tolerance in mice(*53*), and *Smarca5*, a key regulator in DNA replication and DNA damage repair(*54*) (Fig. 4F). These HFD-enriched regions were associated with distinct GO terms related to cellular stress response (Fig. 4G) and composed of genomic regions containing ELK1 (103 sites), ETS2 (151 sites), ELF2 (219 sites), and GABPA (797 sites) motifs (Fig. 4H).

**Fig. 4.**
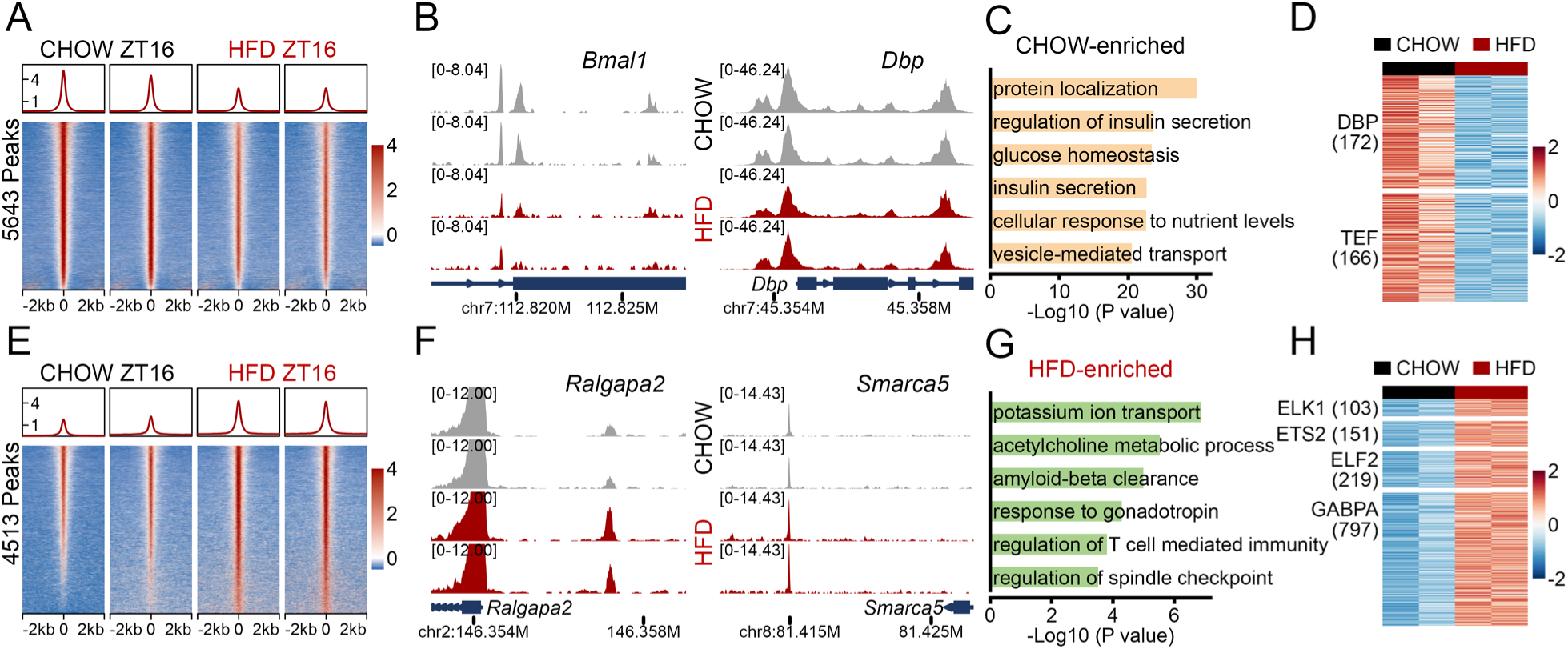
HFD feeding alters chromatin accessibilities of male mice. (A-D) Chromatin regions preferentially accessible in chow-fed mice at ZT16 (5,643 peaks, FDR < 0.1). Average signal profile (top) and heatmaps (bottom) of ATAC-seq signal in each diet (A), with the color scale indicating z-scored chromatin accessibility across a 4-kb window centered on peak summits (−2 kb to +2 kb; 25-bp bins), normalized across all four samples; genome browser tracks at the *Bmal1* and *Dbp* loci (B), unit: CPM; Gene Ontology enrichment for genes associated with these peaks (C); and ATAC-seq signal at the subset containing DBP (172 sites) and TEF (166 sites) motifs among CHOW-enriched peaks (D), with the color scale indicating z-scored chromatin accessibility at motif-containing peaks, normalized across all four samples. (E-H) Regions preferentially accessible in HFD-fed mice at ZT16 (4,513 peaks, FDR < 0.1), presented as in A-D, with browser tracks at the *Ralgapa2* and *Smarca5* loci (F), Gene Ontology enrichment (G) and the motif-containing subset defined by ELK1, ETS2, ELF2, and GABPA motifs among HFD-enriched peaks (H). n=2 mice per time point per group.

We also performed ATAC-seq on male islets at ZT4 (light phase), showing a qualitatively similar but larger-magnitude pattern, with 23,270 chow-enriched peaks (Fig. S4A-D), containing DBP (541 sites)/TEF (535 sites) motifs, and 22,799 HFD-enriched peaks (Fig. S4E-H), containing ELK1 (732 sites)/ETS2 (1,198 sites)/ELF2 (1,403 sites)/GABPA (4,737 sites) motifs, respectively, exemplified by the core clock genes *Bmal1* and *Dbp*, the genes *Ralgapa2* and *Smarca5*. The larger number of differential peaks at ZT4 relative to ZT16 suggests that HFD-induced chromatin remodeling in male islets is most pronounced during the light circadian phase. Collectively, these data show that HFD produces coordinated, time-of-day-dependent shifts in chromatin accessibility at D-box- and ETS-factor-regulated loci, providing a chromatin-level mechanism that parallels and likely underlies the transcriptional changes described above.

### Dark-phase time-restricted feeding rescues circadian β-cell physiology

Having established that HFD disrupts circadian feeding behavior, glucose homeostasis, and islet gene regulation in a coordinated, male-specific manner, we next asked whether directly correcting the disrupted fasting/feeding cycle, independent of total caloric intake or diet composition, could reverse these effects. To test this, male mice on HFD were assigned to one of two feeding regimens: dark phase-restricted feeding (HFDtRFDark, food available only during ZT12-20, the standard active phase), or light phase-restricted feeding (HFDtRFLight, food available only during ZT0-8, the standard inactive phase), followed by behavioral, physiological, and molecular profiling identical to that used above (Fig. 5A). Both time-restricted feeding regimens significantly attenuated HFD-induced weight gain relative to *ad libitum* feeding, with light-phase tRF mice gaining marginally more weight than dark-phase tRF mice by the end of the 8-week protocol (p < 0.05) (Fig. 5B). Both tRF regimens also reduced HFD-induced increases in liver and eWAT (p < 0.05), and lowered blood glucose and plasma insulin relative to *ad libitum* HFD feeding across the 24-hour circadian cycle (p < 0.05) (Fig. S5A-E). By design, tRF reorganized daily activity and food intake rhythms according to the imposed feeding window in each group (Fig. 5C-F). It is important to note that only light-phase tRF group demonstrated a significant decrease in total daily caloric intake, and no differences in total activity were detected among the groups (Fig. 5D, F).

**Fig. 5.**
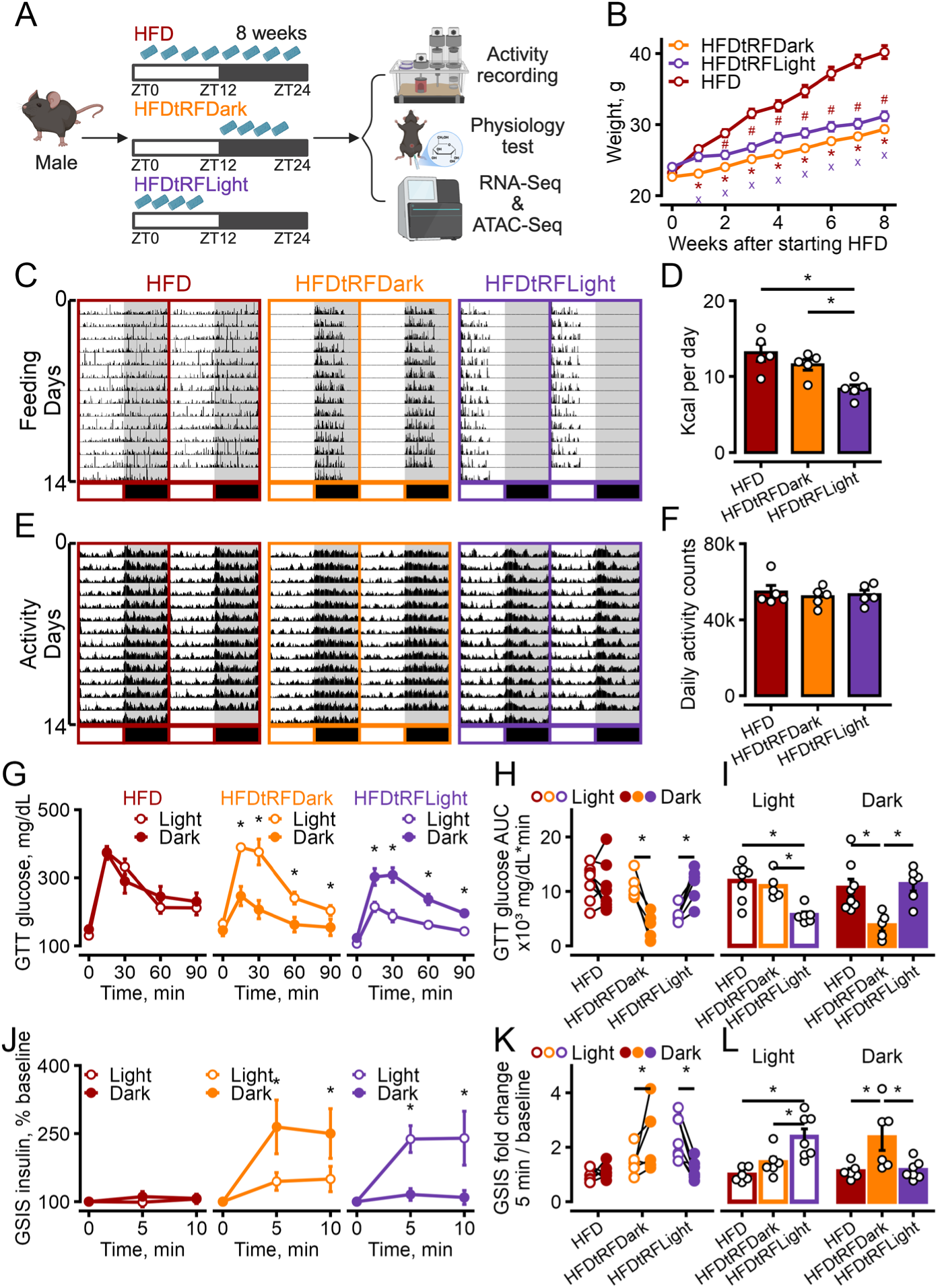
Dark-phase time-restricted feeding (tRF) feeding prevents HFD-induced circadian disruption in glucose tolerance and glucose-stimulated insulin secretion in male mice. (A) Schematic of the feeding-schedule design. HFD-fed male mice were given food *ad libitum*, restricted to the dark phase (HFDtRFDark, ZT12-20), or restricted to the light phase (HFDtRFLight, ZT0-8), followed by feeding and locomotor activity recording, physiology testing, and islet transcriptome and chromatin accessibility profiling. (B) Body weight over 8 weeks in the three groups. n=18 mice for HFD; n=17 mice for HFDtRFDark; n=18 mice for HFDtRFLight. (C, D) Double-plotted feeding actograms, averaged over 5 mice (C) and total daily caloric intake (D). (E, F) Double-plotted locomotor activity actograms, averaged over 5 mice (E) and total daily activity counts (F). n=5 mice per group. White and black bars below feedograms/actograms indicate the light and dark phases of the circadian cycle, respectively. (G-I) GTT. Blood glucose during a glucose tolerance test performed in the light or dark phase in HFD, HFDtRFDark, and HFDtRFLight mice (G); glucose area under the curve (AUC) by light/dark phase within each group (H); and AUC compared between groups within each phase (I). n=8 mice for HFD; n=5 for HFDtRFDark; n=6 for HFDtRFLight. (J-L) GSIS. Plasma insulin (% of baseline) after intraperitoneal glucose administration in the light or dark phase in HFD, HFDtRFDark, and HFDtRFLight mice (J); fold change in insulin secretion (5 min relative to baseline) by light/dark phase within each group (K); and fold change compared between groups within each phase (L). n=6 mice for HFD; n=6 for HFDtRFDark; n=7 for HFDtRFLight. All results were compared by two-way ANOVA with Fisher’s LSD test for post-hoc comparisons. Data are mean ± SEM; * ^# x^ p < 0.05. Specifically, in (B), * Indicates HFD vs. HFDtRFDark; ^#^ indicates HFD vs. HFDtRFLight; ^x^ indicates HFDtRFDark vs. HFDtRFLight.

Dark-phase tRF restored the light/dark difference in glucose tolerance (p < 0.05) that was lost under *ad libitum* HFD feeding, essentially normalizing circadian glucose handling toward the chow-like pattern (Fig. 5G-I). Light-phase tRF, by contrast, produced a robust light/dark difference in glucose tolerance (p < 0.05) that was inverted relative to the normal, chow-like pattern, selectively enhancing glucose tolerance during the light (normally inactive) phase while worsening it during the dark (normally active) phase, effectively creating a new, inverted circadian pattern of glucose tolerance and β-cell function (Fig. 5G-I). GSIS data showed a corresponding pattern: dark-phase tRF restored the circadian rhythm of insulin secretion (p < 0.05) and its fold-change response toward the chow-like state (Fig. 5J-L), whereas light-phase tRF instead produced an inverted rhythm, markedly elevating light-phase insulin secretion above dark-phase levels (p < 0.05) rather than restoring the normal circadian pattern (Fig. 5J-L). In addition, dark-phase tRF significantly improved insulin sensitivity relative to *ad libitum* HFD across both the light and dark phases (p < 0.05), whereas light-phase tRF significantly improved insulin sensitivity relative to *ad libitum* HFD only during the dark phase (p < 0.05) (Fig. S5F-J), indicating that dark-phase feeding confers a more consistent benefit to systemic insulin sensitivity across the day than day-phase feeding. Taken together, these physiological data indicate that aligning feeding to the dark circadian phase is sufficient to restore circadian β-cell physiology disrupted by HFD, whereas feeding during the light phase instead inverts the phase of β-cell circadian rhythmicity, enhancing function specifically in the light phase rather than restoring the normal, chow-like circadian pattern.

### Dark-phase time-restricted feeding partially restores the circadian islet transcriptional program

To determine whether the physiological rescue by dark-phase tRF was accompanied by restoration of the underlying circadian transcriptional program, we performed circadian RNA-seq analysis on islets from both tRF groups (Fig. 6). Comparing overall rhythmic gene set sizes across groups revealed a marked divergence between dark-and light-phase tRF: the number of circadian-rhythmic genes rose from 1,842 in HFD to 4,054 in dark-phase tRF, whereas light-phase tRF instead reduced the rhythmic gene set further, to 713 genes (rhythmicity cutoff: FDR < 0.1) (Fig. 6A-C), indicating that dark-phase tRF, but not day-phase tRF, broadly restores circadian transcriptional rhythmicity in the islet. This same divergence was evident in the specific genes involved: dark-phase tRF promoted *de novo* rhythmicity in 3,472 genes relative to *ad libitum* HFD (Fig. 6D), including the insulin-secretion related gene *Adcy9* (Fig. 6E); this gene set was enriched for insulin secretion and glucose homeostasis related pathways (Fig. 6F), with the amplitude of insulin-secretion genes shifting back toward the chow-like pattern relative to HFD (Fig. 6G). In stark contrast, light-phase tRF enhanced rhythmicity in a considerably smaller set of 593 genes (Fig. 6H), exemplified by *Tomm70a* (Fig. 6I), which was enriched for a functionally diverse set of pathways, most notably “fatty acid metabolic process” (Fig. 6J), with corresponding phase and amplitude changes in this fatty-acid-associated gene set shown in Fig. 6K. This striking difference in both the magnitude and functional identity of the transcriptional response indicates that dark- and light-phase tRF engage largely distinct gene-regulatory programs, rather than producing a shared rescue effect. At the transcription factor motif level, dark-phase tRF promoted a restoration of motif enrichment of chow-associated D-box factors DBP and TEF with 29.9% of their target genes, which lost rhythmicity in HFD, regaining rhythmicity under dark-phase tRF compared to HFD (Fig. 6L, M). In contrast, light-phase tRF produced a bimodal-pattern of motif enrichment for DBP/TEF, enriched at ZT4 and ZT12 (Fig. 6L), with only 7.5% of their target genes, which became arrhythmic in HFD, regaining rhythmicity under light-phase tRF compared to HFD (Fig. 6M). Regarding the HFD-enriched ELK1/ETS2/ELF2/GABPA motifs, both dark-phase and light-phase tRF drove a reduction of this motif activity and the corresponding fraction of their target genes compared to HFD (Fig. 6N, O). Together, these results indicate that night-phase tRF specifically re-engages the D-box-driven insulin-secretion transcriptional program that is disrupted by HFD, whereas day-phase tRF instead induces a distinct rhythmic response that does not reengage normal islet circadian gene expression.

**Fig. 6.**
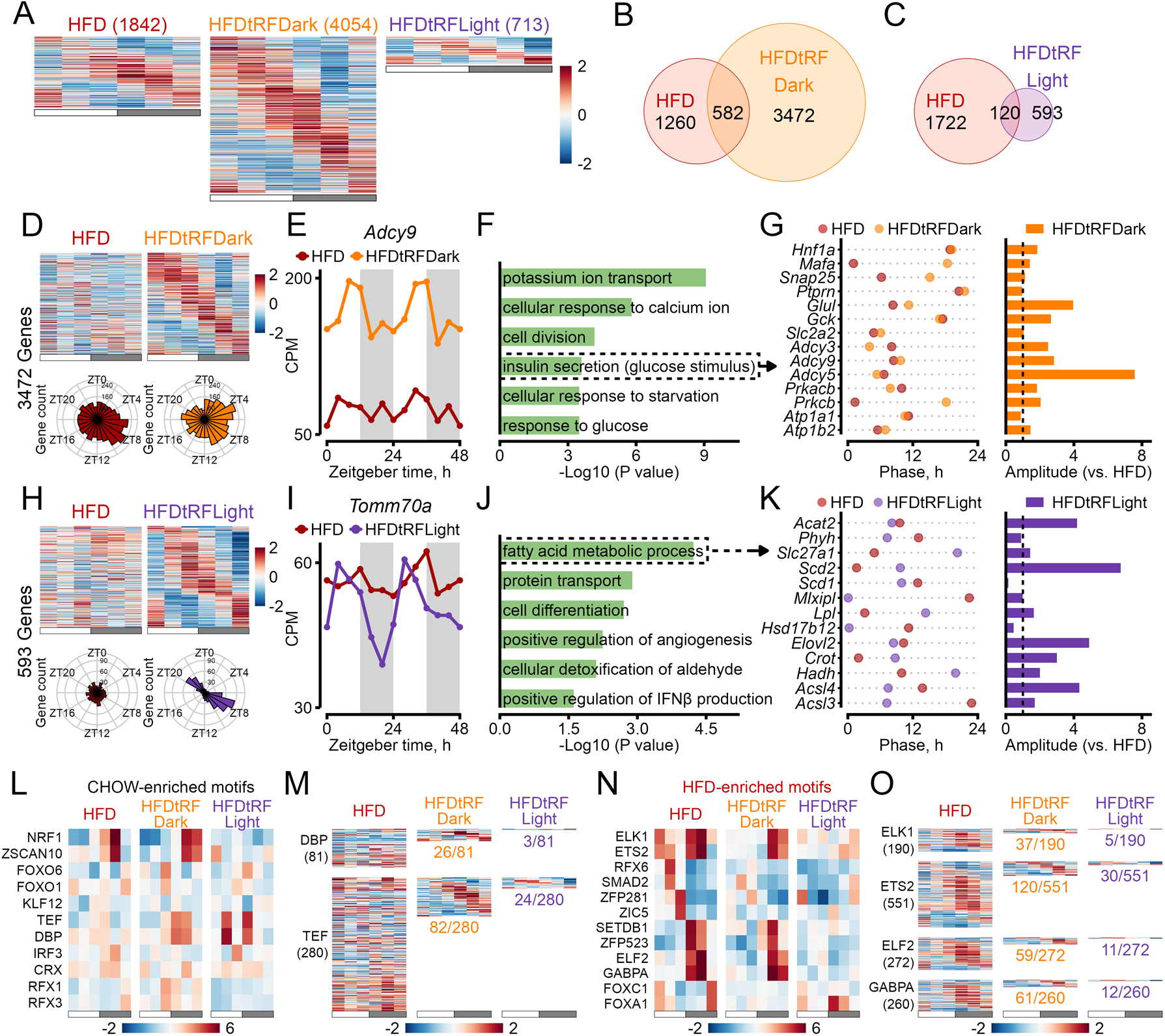
Dark-phase time-restricted feeding partially restores the circadian islet transcriptional program. (A) Heatmaps of expression across the 24-h cycle for genes rhythmic in HFD (1,842 genes, FDR < 0.1), HFDtRFDark (4,054 genes, FDR < 0.1), and HFDtRFLight (713 genes, FDR < 0.1) mice. Columns represent Zeitgeber time (ZT) points sampled every 4 h (from ZT0 to ZT20), with values averaged from two replicates (mice). Gene expression was z-score normalized across HFD, HFDtRFDark, and HFDtRFLight groups. Genes are ranked by their peak-expression phase. White and gray bars below the heatmaps indicate the light and dark phases of the circadian cycle, respectively. (B, C) Venn diagrams of the HFD gene set with HFDtRFDark (B; 582 shared) and with HFDtRFLight (C; 120 shared). (D-G) Genes with *de novo* rhythmicity in HFDtRFDark relative to HFD (3,472 genes). Heatmaps of expression across the 24-h cycle with polar plots of peak-phase distribution in each group (D); expression across Zeitgeber time for the representative gene *Adcy9* (E); Gene Ontology enrichment for the cluster, with “insulin secretion involved in cellular response to glucose stimulus” highlighted, shorten as “insulin secretion (glucose stimulus)” on the graph (F); and peak phase and amplitude (G) of insulin-secretion genes in HFDtRFDark relative to HFD. (H-K) Genes with *de novo* rhythmicity in HFDtRFLight relative to HFD (593 genes), presented as in D-G, with the representative gene *Tomm70a* (I) and “fatty acid metabolic process” highlighted (J, K). For (E, I), data from the two replicates per time point were plotted sequentially over a 48-h period, with replicate 1s from each time point displayed from 0-20 h and replicate 2s displayed from 24-44 h. The data point at 48 h is identical to that at 0 h and is included to facilitate visualization of circadian rhythmicity. (L, M) Enrichment of chow-enriched transcription factor motifs in rhythmic genes across HFD, HFDtRFDark, and HFDtRFLight (L); and expression across the 24-h cycle of DBP and TEF ChIP-seq target genes that lose rhythmicity under HFD compared to chow, with the proportion regaining rhythmicity indicated for each tRF group (M). (N, O) Enrichment of HFD-enriched transcription factor motifs in rhythmic genes across HFD, HFDtRFDark, and HFDtRFLight (N); ELK1, ETS2, ELF2, and GABPA ChIP-seq target genes that gain rhythmicity under HFD compared to chow, with the proportion still be rhythmic indicated for each tRF group (O). Color scales in (A, D, H, M, and O) depict z-scored gene expression, whereas those in (L and N) depict normalized enrichment scores (NES) from RcisTarget.

### Dark-phase time-restricted feeding redirects islet chromatin accessibility toward a chow-like state

Finally, to determine whether the transcriptional differences between dark-and light-phase tRF were mirrored at the level of chromatin accessibility, we performed ATAC-seq on islets from both tRF groups at ZT4 and ZT16. At ZT16, dark-phase tRF resulted in loss of 4,974 HFD-associated peaks (Fig. 7A, B) and a gain of 5,109 peaks (Fig. 7E, F) relative to *ad libitum* HFD (FDR < 0.1), whereas light-phase tRF produced a qualitatively distinct reorganization, with 3,362 peaks lost (Fig. 7C, D) and 6,617 peaks gained (Fig. 7G, H) (FDR < 0.1). Exemplified chromatin regions are shown on Fig. 7I and K. GO term enrichment showed that the HFD-associated peaks lost with either regimen were dominated by cell-adhesion and inflammatory/immune-related processes, including cadherin-mediated cell-cell adhesion, chemotaxis, and I-κB phosphorylation (Fig. 7B, D), consistent with the NF-κB-driven inflammatory/stress program identified under *ad libitum* HFD conditions (Fig. 3L-O, Fig. S3B). In contrast, the peaks gained with dark-phase tRF were enriched for metabolic processes, including insulin secretion, glucose homeostasis, and response to glucose (Fig. 7F), whereas peaks gained with light-phase tRF were enriched for response to glucose, but also negative regulation of insulin secretion (Fig. 7H). Consistent with this, dark-phase tRF produced a markedly stronger restoration of DBP (172 sites) and TEF (166 sites) motif accessibility at ZT16 than light-phase tRF, which showed only a weak, partial increase relative to HFD (Fig. 7J). Both dark-phase tRF and day-phase tRF promoted a strong reduction of HFD-induced ELK1/ETS2/ELF2/GABPA motif accessibility (Fig. 7L).

**Fig. 7.**
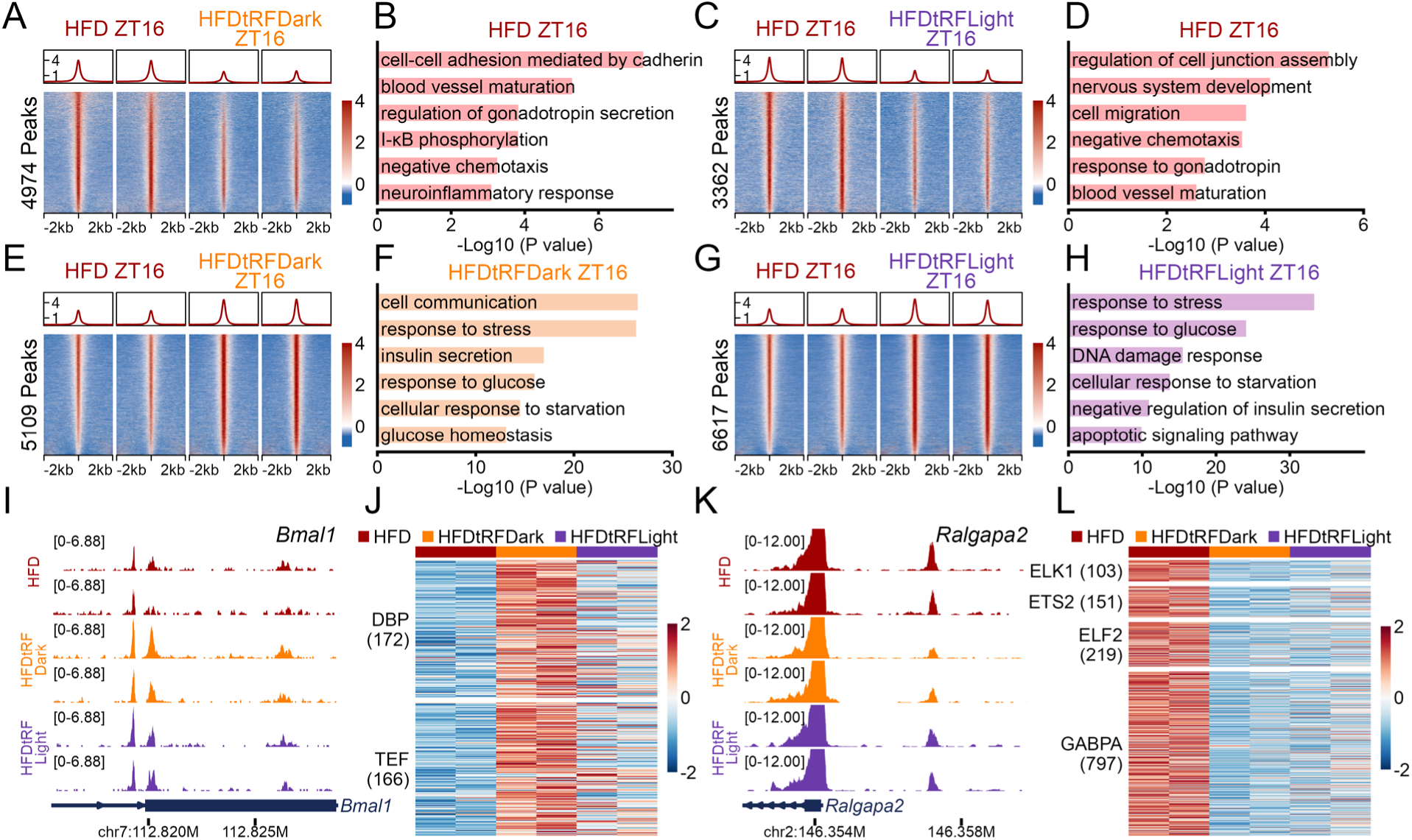
Dark-phase tRF promotes chromatin accessibility of genes important in the regulation of β-cell function. (A-H) Differentially accessible regions at ZT16. Average signal profile (top) and heatmaps (bottom) of ATAC-seq signal and Gene Ontology enrichment are shown for regions having reduced accessibility with dark-phase tRF (4,974 peaks, FDR < 0.1; A, B) or with light-phase tRF (3,362 peaks, FDR < 0.1; C, D), and for regions having increased accessibility with dark-phase tRF (5,109 peaks, FDR < 0.1; E, F) or with light-phase tRF (6,617 peaks, FDR < 0.1; G, H), each relative to *ad libitum* HFD. (I-L) Accessibility of representative genomic loci and motif-containing regions at ZT16 across HFD, HFDtRFDark, and HFDtRFLight conditions. Genome browser tracks at the *Bmal1* locus (I, unit: CPM) and ATAC-seq signal at regions containing DBP (172 sites) and TEF (166 sites) motifs that lose accessibility under HFD compared to chow (J). Genome browser tracks at the *Ralgapa2* locus (K, unit: CPM) and ATAC-seq signal at regions containing ETS-family motifs that gain accessibility under HFD compared to chow (L). n=2 mice per time point per group. Color scales in (A, C, E, G) indicate z-scored chromatin accessibility across a 4-kb window centered on peak summits (−2 kb to +2 kb; 25-bp bins), normalized across all samples shown in each comparison. Color scales in (J, L) indicate z-scored chromatin accessibility at motif-containing regions, normalized across HFD, HFDtRFDark, and HFDtRFLight samples.

A qualitatively similar, though larger-magnitude, pattern was observed at ZT4, with dark- and light-phase tRF producing distinct sets of lost (17,331 and 24,737 peaks, respectively; Fig. S6A-D) and gained (19,350 and 26,842 peaks, respectively; Fig. 7E-H) peaks (FDR < 0.1), exemplified by *Dbp* and *Smarca5* (Fig. S6I, K). As at ZT16, the HFD-associated peaks lost with either regimen were enriched for cell-adhesion and inflammatory-related terms, including I-κB phosphorylation and chemokine production (Fig. S6B, D). The peaks gained with dark-phase tRF remained enriched for insulin secretion, glucose homeostasis, and also response to stress (Fig. S6F); peaks gained with light-phase tRF were enriched for more cellular stress and apoptotic pathways in addition to the insulin secretion (Fig. S6H), directly opposing the beneficial metabolic signature seen with dark-phase tRF. Both dark-phase and light-phase tRF partially restored DBP/TEF motif accessibility (Fig. S6J), and reduced HFD-induced ELK1/ETS2/ELF2/GABPA motif accessibility at ZT4 time point (Fig. S6L). Collectively, these chromatin-level findings closely parallel the transcriptional results described above, indicating that dark-phase, but not light-phase, tRF redirects chromatin accessibility toward the D-box-regulated, chow-like regulatory state at genes governing key β-cell functions, providing a chromatin-level basis for the selective physiological rescue observed with dark-phase feeding.

## Discussion

Together, our findings identify a feeding-rhythm-dependent mechanism by which DIO disrupts circadian β-cell physiology in a sex-specific manner. In male mice, HFD feeding blunts the normal day-night feeding rhythm and this feeding-rhythm-specific disruption is accompanied by 1) a coordinated loss of circadian glucose tolerance and glucose-stimulated insulin secretion, 2) reorganization of the circadian islet transcriptome and chromatin landscape, and 3) a shift in the dominant circadian transcriptional regulators from D-box factors (DBP, TEF) toward ETS-family factors (ELK1, ETS2, ELF2, GABPA). Notably, the origin of this feeding-rhythm disruption by HFD has recently been traced to a vagus nerve-mediated liver-to-brain circuit(*29*). Female mice, in contrast, largely preserve their feeding rhythm and circadian glucose homeostasis despite HFD-induced weight gain, suggesting that the metabolic consequences of DIO on the circadian islet are not exclusively a function of adiposity but also track closely with the degree of feeding-rhythm disruption.

Mechanistically, our data suggests that HFD does not merely dampen the islet circadian clock but actively reprograms its transcriptional output. The loss of rhythmicity in the D-box-driven, insulin-secretion-associated gene program occurs alongside the emergence of a distinct, ETS-family-associated inflammatory/stress rhythmic program that is largely absent in the chow-fed state. This is consistent with previous reports that HFD disrupts behavioral and molecular circadian rhythms broadly(*28*), and extends that concept by showing that, at the islet level, this disruption is not simply a flattening of existing rhythms but a qualitative shift in which genes and pathways are placed under circadian control. The persistence of a core clock gene cluster (including *Dbp* itself) with reduced but not abolished amplitude further suggests that the circadian oscillator in the islet remains intact under DIO, and that the downstream, tissue-specific outputs of that clock are the primary site of dysfunction.

Evidence from other metabolic tissues, including liver, muscle, and adipose tissues, indicates that high-fat feeding does not simply dampen the core oscillator but instead reprograms circadian output. In mouse liver, HFD induces extensive clock remodeling and generates large numbers of *de novo* oscillating transcripts through two convergent mechanisms: impaired CLOCK: BMAL1 recruitment to chromatin and cyclic activation of alternative transcriptional pathways via PPARγ(*30*). At the enhancer level, DIO drives widespread remodeling of circadian enhancer activity, imposing synchronous, high-amplitude rhythms of both fatty acid synthesis and oxidation through rhythmic activation of SREBP and PPARα(*55*). These effects extend beyond hepatocytes. Obesity and insulin resistance have been shown to reprogram BMAL1 activity in liver, muscle, and adipose tissue, and these effects have been linked to systemic insulin resistance since corresponding treatment with a known insulin sensitizer, rosiglitazone, reversed HFD-induced changes in BMAL1 chromatin recruitment(*56*). Comparable obesity-driven redistribution has also been reported for the diurnal glucocorticoid response(*57*). Collectively, these findings support a model of DIO-induced cistrome plasticity rather than clock failure. In addition, the enrichment of ETS2, ELK1, ELF2, and GABPA motifs under HFD suggest different routes through which HFD could reshape circadian regulation in pancreatic islets. GABPA controls the majority of nuclear-encoded OXPHOS subunits, mito/ribosomal proteins, and factors required for mitochondrial DNA replication and transcription(*58*), placing it directly within the mitochondrial-capacity pathway impacted by HFD. In contrast, transcriptional activities of ELK1 and ETS2 are regulated by MAPK/ERK-mediated phosphorylation(*59–61*), and the ERK signaling has been shown to be activated by insulin via the Ras–RAF–MEK–ERK pathway(*62*). We previously reported that hyperinsulinemia represses DBP expression by inhibiting FOXO1(*10*). Together with the data here, this suggests that HFD-associated hyperinsulinemia might be the underlying driver that promotes the shift of circadian regulators from DBP to ETS-family factors: enhancing ETS-dependent transcription via ERK signaling while suppressing DBP-dependent output through FOXO1 inhibition.

The sexual dimorphism we observe fits within a growing recognition that circadian physiology differs systematically between the sexes. In humans, the intrinsic period of the circadian pacemaker is shorter in women than in men(*41*), and women showed an earlier phase of core body temperature and melatonin rhythms compared with men relative to sleep onset(*43*). Analysis of post-mortem human tissue has shown that the organization of 24-hour transcriptional rhythms is itself sex-dimorphic and diverges further with age, with roughly twice as many rhythmic genes overall detected in females as in males across all amplitude thresholds(*39*). Rodent studies point in the same direction: female mice show a larger rhythmic fraction of the liver transcriptome and metabolome than males(*63, 64*). Female mice are also more resilient to chronic circadian misalignment: under a shiftwork-mimicking protocol of weekly 8-hour phase advances on a high-fat diet, females preserved behavioral and transcriptional rhythmicity and were protected from the cardiometabolic consequences observed in males, a sex difference that extended to activity rhythmicity and metabolic syndrome incidence among human shiftworkers(*65*). Our results extend this pattern into circadian feeding rhythm and pancreatic islets, where female mice preserved rhythmic islet function and feeding behavior under a dietary challenge sufficient to disrupt both in males.

Our time-restricted feeding experiments show that this male-specific circadian dysfunction is reversible, but only when feeding is restricted to the active (dark) phase. Dark-phase tRF, independent of any change in caloric intake or diet composition, restored circadian glucose tolerance and GSIS, broadly restored the circadian islet transcriptome, and partially re-established D-box transcription factor activity, consistent with previous work showing that time-restricted feeding can protect against circadian disruption of β-cell function through DBP activation(*10*). Notably, light-phase tRF did not simply fail to rescue the HFD phenotype; it actively created a new, inverted pattern of circadian β-cell physiology, with enhanced glucose tolerance and insulin secretion specifically during the light (inactive) phase, and a transcriptional and chromatin-accessibility signature dominated by cellular stress and interferon-associated programs. This finding underscores that the timing of time-restricted feeding relative to the endogenous active phase, and not simply the imposition of a fasting/feeding cycle, determines whether the intervention is beneficial or potentially maladaptive.

While our findings were obtained in a mouse model of DIO, they carry a specific translational implication: the effectiveness of time-restricted eating may depend not only on the duration of the feeding window but on its placement relative to the endogenous circadian phase. The premise that such a feeding window can reposition peripheral circadian phase in humans is established: a 5-h meal delay shifts plasma glucose and adipose PER2 rhythms without altering melatonin or cortisol(*66*), demonstrating that feeding time entrains peripheral tissue independently of the central pacemaker, the necessary condition for the inverted phenotype we observe in mice. Consistent with this, experimental circadian misalignment in humans reduces glucose tolerance and impairs early-phase insulin secretion(*67*). Conversely, when the feeding window is aligned with the active phase, early time-restricted eating improves insulin sensitivity, β-cell responsiveness and blood pressure without weight loss(*68*) and outperforms a mid-day window for insulin sensitivity(*69*). Taken together with our data, this pattern suggests that the important variable is neither the imposition of a fasting/feeding cycle nor phase shifting itself, but the position of the feeding window relative to the endogenous active phase, with alignment restoring rhythmic β-cell function, whereas anti-phase feeding generates a distinct, stress- and interferon-associated program.

Collectively, our findings establish feeding-rhythm disruption as a proximal, sex-specific driver of circadian β-cell dysfunction in DIO and identify the alignment of feeding time with the endogenous active phase, rather than time-restricted feeding itself, as the critical determinant of whether such an intervention restores or further disrupts circadian transcriptome, chromatin accessibility, and metabolic health. At the genomics level, our data links DIO induced circadian alteration to a reorganization of the islet regulatory landscape from D-box– driven to ETS-type transcriptional control, nominating DBP/TEF and ETS-family factors as candidate effectors through which feeding time is translated into rhythmic islet gene expression.

## Materials and Methods

### Animals and experimental study design

All animal procedures were approved by the Mayo Clinic Institutional Animal Care and Use Committee. In total, this study used 122 male and 80 female C57BL/6J mice (the Jackson Laboratory, Bar Harbor, ME). All mice were housed in the Promethion metabolic cage system (Sable Systems International, Las Vegas, NV) within external light-proof cabinets (Phenome Technologies, Lincolnshire, IL) and fed standard chow diet (LabDiet, St. Louis, MO, Cat# 5053) or high-fat diet (Research Diets, New Brunswick, NJ, Cat# D12492). By convention, ZT0 was denoted as the onset of the light cycle, and ZT12 was denoted as the onset of the dark cycles. During an initial 2-week acclimatization period, all animals were entrained to a 12-hour light/12-hour dark cycle with ad libitum access to water and standard chow diet. Following the 2-week period, the animals were assigned to 4 different experimental groups for 8 weeks: (i) CHOW group: ad libitum access to water and chow diet; (ii) HFD group: ad libitum access to water and high-fat diet; (iii) HFDtRFDark group: ad libitum access to water with high-fat diet food access only between ZT12 and ZT20; (iv) HFDtRFLight group: ad libitum access to water with high-fat diet food access only between ZT0 and ZT8. Temporal access to feeding hopper was controlled by MetaScreen v2.3.15 (Sable Systems International), which allowed remote/noninvasive opening and closing of the feeding trough in individual mouse cages.

### Assessment of circadian food intake and motor activity

Circadian rhythms in feeding and activity were assessed in mice individually housed in Promethion metabolic cages outfitted with a Promethion MM-1 food intake load cell (at 3-mg resolution) and BXY beambreak activity monitor system to track gross motor activity. Raw data of food intake and activity was processed and binned into 1-min intervals by MacroInterpreter v23.6 (Sable Systems International). Feedograms and actograms were subsequentially generated using Feedogram Toolbox (https://zhangluhui.github.io/feedogram_toolbox/) and Actogram Toolbox (https://zhangluhui.github.io/actogram_toolbox/) respectively.

### Assessment of in vivo glucose tolerance, glucose-stimulated insulin secretion, and insulin tolerance

Diurnal in vivo glucose tolerance, glucose-stimulated insulin secretion, and insulin tolerance were assessed in each mouse at both ZT4 and ZT16. For glucose tolerance test, mice were injected in the intraperitoneal space with a bolus of dextrose (2 g/kg of body mass) (Hospira Inc, Lake Forest, IL, NDC# 00409-6648-02). Blood glucose was then sampled and measured by the Accu Chek Guide Me Glucose Meter (Roche, Switzerland) at baseline and subsequently at 15, 30, 60, and 90 min after dextrose injection. For glucose-stimulated insulin secretion, mice were injected in the intraperitoneal space with a bolus of dextrose (2 g/kg of body mass), and blood was then collected at baseline and subsequently at 5 and 10 min after dextrose injection. Blood samples were centrifuged by 2500 g for 15 min at 4°C, and plasma was then collected for insulin measurement via ELISA (ALPCO, Ref# 80-INSMS-E10). For insulin tolerance, mice received a bolus of insulin (0.75 mU/g of body mass) (Novo Nordisk, Plainsboro, NJ, NDC# 0169-1833-11) via intraperitoneal injection, and blood glucose was measured at baseline and subsequently at 15, 30, 60, and 90 min after injection.

### Mouse islet isolation, RNA isolation, and RNA sequencing (RNA-Seq)

Mouse islets were isolated at ZT0, ZT4, ZT8, ZT12, ZT16, and ZT20, with standard collagenase (Roche, Switzerland, Cat# 05401020001) digestion and histopaque (Sigma-Aldrich, St. Louis, MO, Cat# 10771-100ML) gradient centrifugation method. Total RNA was isolated from mouse islets using the RNeasy mini kit (Qiagen, Hilden, Germany, Cat# 74160). RNA was then shipped on dry ice to Genewiz (South Plainfield, NJ) for quality assessment and sequencing. Specifically, RNA size and quality were assessed using the Tape Station Bionalyzer (Agilent, Santa Clara, CA), and concentration was determined by Qubit fluorometry. RNA-Seq libraries were prepared with the NEBNext Ultra II RNA Library Prep Kit for Illumina (New England Biolabs, Ipswich, MA) using the polyA selection method. >25 million paired-end, 150 base pair (bp) reads per sample were then sequenced on an Illumina HiSeq instrument.

For RNA-Seq data processing, the raw reads were mapped onto the mouse reference genome mm39, using STAR v2.7.9a(*70*) with the setting “--quantMode GeneCounts” and the genome annotation from NCBI RefSeq to obtain gene counts. The raw counts were then normalized by the R package edgeR v4.8.2(*71*) using the default Trimmed Mean of M-values (TMM) method, outputting the counts per million (CPM) values. Genes that were expressed at a level at least 1 CPM in at least 2 samples were included in downstream analysis. To identify circadian transcripts in mouse islet samples, normalized count matrices were imported into MetaCycle v1.2.1(*49*) and circadian rhythmicity was determined by ARSER method(*48*) with default settings. Genes were considered significantly rhythmic at a false discovery rate (FDR) < 0.1. Significantly rhythmic genes were subjected to biological pathway enrichment using the Database for Annotation, Visualization, and Integrated Discovery (DAVID)(*72, 73*) (National Institute of Health, Bethesda, MD). Motif enrichment analysis was performed among rhythmic genes with R package RcisTarget v1.29.0(*50, 74*).

### Mouse islet isolation, and Assay for Transposase-Accessible Chromatin using sequencing (ATAC-seq)

For bulk ATAC-seq, mouse islets were freshly isolated at ZT4 and ZT16 using the same isolation method described above. Specifically, after being washed with ice-cold PBS, islets were incubated in the lysis buffer (1% Igepal CA-630, 3 mM MgCl2, 10 mM NaCl, 10 mM Tris buffer) on ice for 25 min. The nuclei were then pelleted down via centrifugation (500 g for 10 min, 4 °C). After resuspension in the lysis buffer, the nuclei were transferred to the Mayo Clinic Epigenomics Development Laboratory (Rochester, MN) for the ATAC-seq library preparation following the omni-ATAC protocol(*75*). The libraries were then shipped on dry ice to Genewiz (South Plainfield, NJ) for quality assessment and sequencing. >50 million paired-end, 150 base pair (bp) reads per sample were sequenced on an Illumina HiSeq instrument.

For ATAC-Seq data processing, sequencing adapters and low-quality sequencing reads were removed using Trimmomatic v0.39(*76*) initially. Reads were then aligned to the mm39 reference genome assembly using Bowtie2 v2.3.3.1(*77*). Picard MarkDuplicates v2.26.10(*78*) was used to identify the duplicated sequencing reads. Mitochondrial reads, duplicated reads, and reads with low mapping scores were then filtered out with Samtools v1.20(*79*). Peak calling for individual samples was performed with MACS3 v3.0.3(*80*). R package Diffbind v3.20.0(*81*) was then utilized for normalization and differential analysis among different samples via the built-in EdgeR(*71*) method, with differential cutoff being FDR < 0.1. Differentially accessible chromatin regions were subjected to biological pathway enrichment analysis using the R package rGREAT v2.14.0(*82*). Software MEME FIMO v5.5.7(*83*) was used to scan for transcription factor motif enrichment among differentially accessible chromatin regions. The genome browser track files (bigWig format) were generated with deepTools bamCoverage v3.5.5(*84*), employing the normalization factor calculated by DiffBind, and the bigWig files were subsequentially visualized via the R package Trackplot v1.6.0(*85*). The peak-centered heatmaps were generated using the R package EnrichedHeatmap v1.42.0(*86*).

### Statistical analysis

Results presented are expressed as the mean ± SEM. Statistical analysis was completed using GraphPad Prism v10.6.1 (Dotmatics Inc., Boston, MA). Statistical analysis was performed by either two-tailed Student’s t test for comparison of two independent groups, one-way analysis of variance (ANOVA) or two-way ANOVA for comparison of three or more groups with Fisher’s LSD test for post-hoc comparisons. The value of p < 0.05 was considered statistically significant, unless otherwise noted.

## Acknowledgments

We thank L. D. Pham for technical assistance with ATAC-Seq experiments. Illustrations of study design and analysis were created with Biorender.com premium service. **Funding**: We acknowledge funding support from the NIH (R01DK098468 to A.V.M.). **Author contributions**: L.Z. contributed to study design, conducted experiments, and assisted with the data analysis, interpretation, and preparation of the manuscript. S.K.S., K.R., T.N., K.O., and A.B. conducted experiments and assisted with the data analysis, interpretation, and preparation of the manuscript. A.G.M. and Z.W. contributed to study design, data analysis, interpretation, and preparation of the manuscript. A.V.M. designed, interpreted the studies, and wrote the manuscript with L.Z. **Competing interests**: The authors declare that they have no competing interests. **Data and materials availability**: All data needed to evaluate the conclusions in the paper is present in the paper and the Supplementary Materials.

