## Supplemental Figures for "Circadian feeding rhythm loss drives sex-specific β-cell dysfunction in diet-induced obesity"

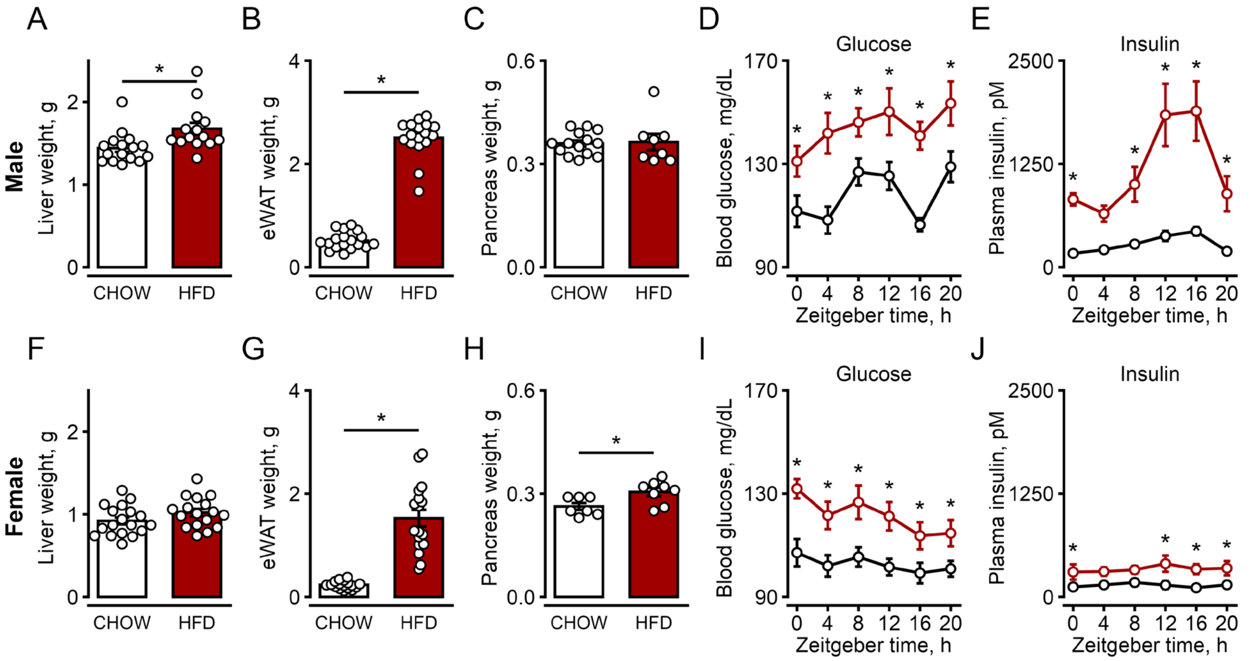


**Fig. S1. HFD feeding promotes accumulation of epididymal white adipose (eWAT), hyperglycemia, hyperinsulinemia in both male and female mice.** (A-E) Male mice. Liver (n=16 mice for CHOW; n=14 mice for HFD) (A), epididymal white adipose tissue (n=18 mice for CHOW; n=17 mice for HFD) (eWAT; B), and pancreas (n=14 mice for CHOW; n=8 mice for HFD) (C) weights after 8 weeks of chow or HFD feeding; blood glucose (n=9 mice per ZT time point) (D) and plasma insulin (n=8 mice per ZT time point, except n=6 mice for HFD ZT0, 4, and 20) (E) measured across the 24-h cycle. (F-J) Female mice, presented as in a-e. Liver: n=18 mice for CHOW; n=17 mice for HFD. eWAT: n=17 mice for CHOW; n=17 mice for HFD. For pancreas: n=7 mice for CHOW; n=8 mice for HFD. For (A-C, F-H), results were compared by unpaired two-tailed Student's t test. For (D, E, I, J), results were compared by two-way ANOVA with Fisher's LSD test for post-hoc comparisons. Data are mean ± SEM; $*p < 0.05$.

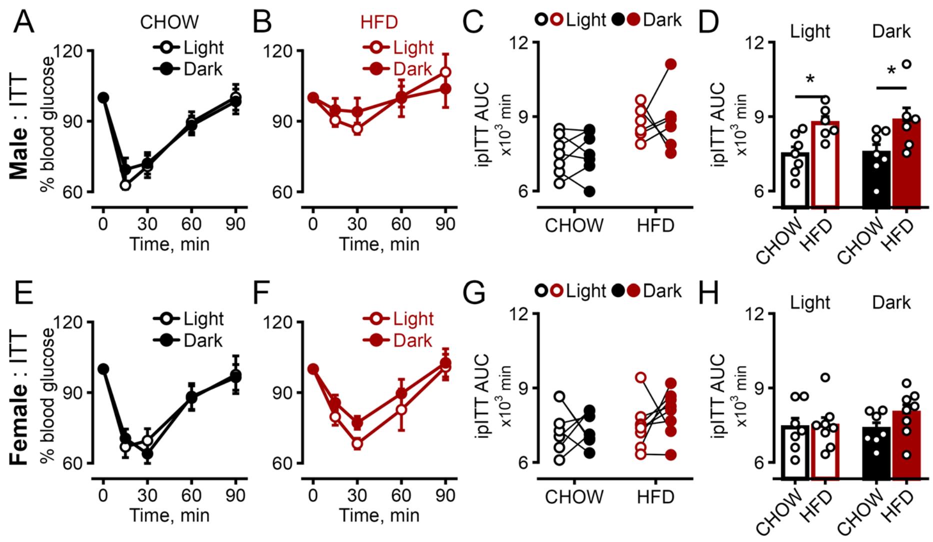


**Fig. S2. HFD feeding promotes glucose intolerance in male mice but not in females.** (A-D) Male mice. Blood glucose during an intraperitoneal insulin tolerance test (ipITT) performed in the light or dark phase in chow-fed (n=7 mice) (A) and HFD-fed (n=6 mice) (B) mice; glucose area under the curve (AUC) by light/dark phase within each diet (C); and AUC compared between diets within each phase (D). (E-H) Female mice, presented as in A-D. n=7 mice for CHOW; n=8 mice for HFD. All results were compared by two-way ANOVA with Fisher's LSD test for post-hoc comparisons. Data are mean ± SEM; *p < 0.05.

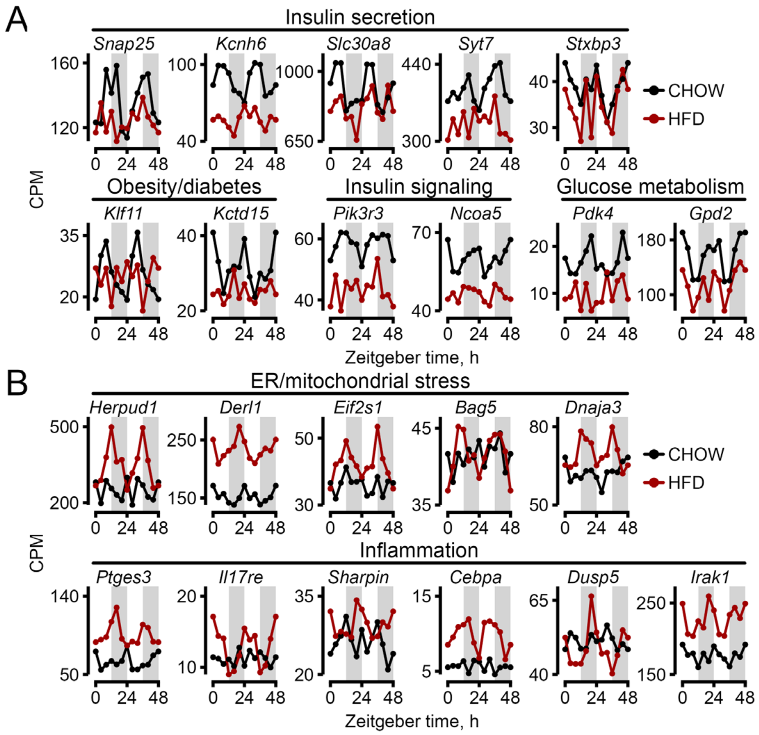


**Fig. S3. Representative circadian transcripts altered by HFD in pancreatic islets.** (A) Temporal expression profiles of DBP- and TEF-target genes involved in insulin secretion, insulin signaling, and glucose metabolism. (B) Temporal expression profiles of ELK1-, ETS2-, ELF2-, and GABPA-target genes associated with ER/mitochondrial stress and inflammation. Gene expression data were generated from islets isolated for RNA-seq at 4-h intervals across the 24-h circadian cycle, with two independent replicates (mice) collected at each time point. To facilitate visualization of rhythmic expression patterns, the two replicates collected at each Zeitgeber time were plotted sequentially over a 48-h interval. Replicate 1 is shown from 0-20 h and replicate 2 from 24-44 h. The 48-h time point is identical to the 0-h time point and was included solely to close the circadian cycle.

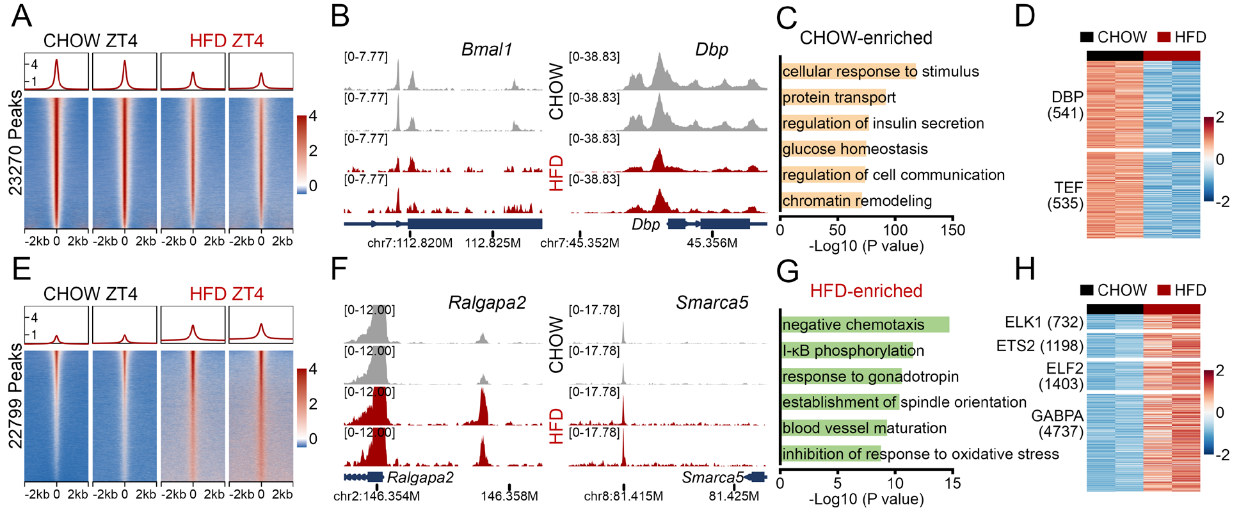


**Fig. S4. HFD feeding alters chromatin accessibilities of male mice at ZT4.** (A-D) Chromatin regions preferentially accessible in chow-fed mice at ZT4 (23,270 peaks, FDR < 0.1). Average signal profile (top) and heatmaps (bottom) of ATAC-seq signal in each diet (A), with the color scale indicating z-scored chromatin accessibility across a 4-kb window centered on peak summits (−2 kb to +2 kb; 25-bp bins), normalized across all four samples; genome browser tracks at the *Dbp* locus (B), unit: CPM; Gene Ontology enrichment for genes associated with these peaks (C); and ATAC-seq signal at the subset containing DBP (541 sites) and TEF (535 sites) motifs among CHOW-enriched peaks (D), with the color scale indicating z-scored chromatin accessibility at motif-containing peaks, normalized across all four samples. (E-H) Regions preferentially accessible in HFD-fed mice at ZT4 (22,799 peaks, FDR < 0.1), presented as in A-D, with browser tracks at the *Smarca5* locus (F) and the motif-containing subset defined by ELK1, ETS2, ELF2, and GABPA motifs among HFD-enriched peaks (H). n=2 mice per time point per group.

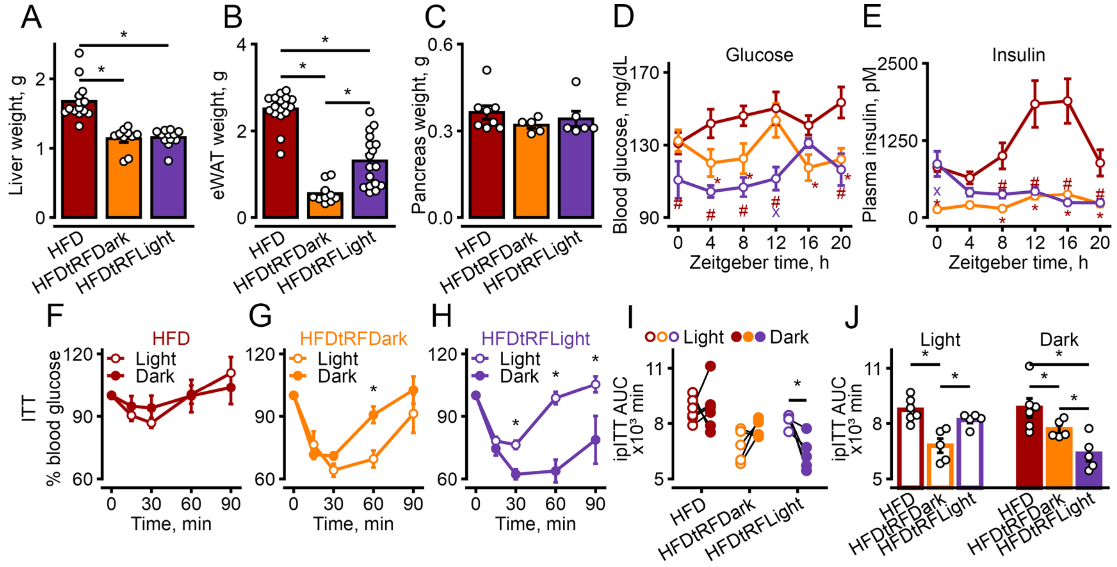

**Fig S5. Both dark- and light-phase tRF inhibit weight gain of liver and epididymal white adipose (eWAT), hyperglycemia, hyperinsulinemia, and enhance insulin sensitivity.** (A-C) Liver (n=14 mice for HFD; n=10 mice for HFDtRFDark; n=12 mice for HFDtRFLight) (A), epididymal white adipose tissue (n=17 mice for HFD; n=10 mice for HFDtRFDark; n=17 mice for HFDtRFLight) (eWAT; B), and pancreas (n=8 mice for HFD; n=5 mice for HFDtRFDark; n=6 mice for HFDtRFLight) (C) weights in HFD, HFDtRFDark, and HFDtRFLight mice. (D, E) Blood glucose (n=9 mice for HFD; n=5 mice for HFDtRFDark; n=6 mice for HFDtRFLight ZT0, 16, 20 and n=7 mice for ZT4, 8, 12) (D) and plasma insulin (n=6 mice for HFD ZT0, 4, 20 and n=8 for ZT8, 12, 16; n=6 mice for HFDtRFDark; n=6 mice for HFDtRFLight ZT0, 16, 20 and n=7 mice for ZT4, 8, 12) (E) measured across the 24-h cycle in the three groups. (F-J) Insulin sensitivity. Blood glucose during an intraperitoneal insulin tolerance test (ipITT) performed in the light or dark phase in HFD (n=6 mice) (F), HFDtRFDark (n=5 mice) (G), and HFDtRFLight (n=5 mice) (H) mice; glucose area under the curve (AUC) by light/dark phase within each group (I); and AUC compared between groups within each phase (J). All results were compared by two-way ANOVA with Fisher's LSD test for post-hoc comparisons. Data are mean ± SEM; $^{*\,\#\,x}p < 0.05$. Specifically, for (D) and (E): * Indicates HFD vs. HFDtRFDark; # indicates HFD vs. HFDtRFLight; x indicates HFDtRFDark vs. HFDtRFLight.

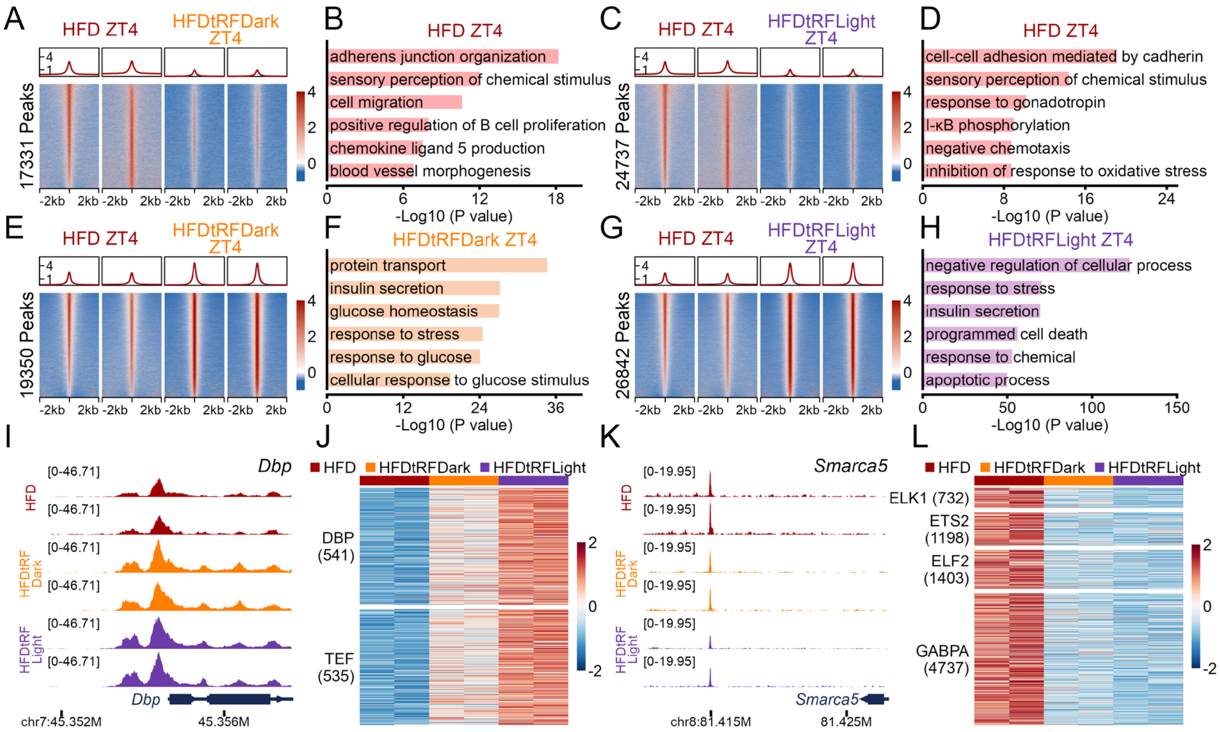


**Fig. S6. Dark-phase tRF promotes chromatin accessibilities of genes related to key β-cell functions at ZT4.** (A-H) Differentially accessible regions at ZT4. Average signal profile (top) and heatmaps (bottom) of ATAC-seq signal and Gene Ontology enrichment are shown for regions having reduced accessibility with dark-phase tRF (17,331 peaks, FDR < 0.1; A, B) or with light-phase tRF (24,737 peaks, FDR < 0.1; C, D), and for regions having increased accessibility with dark-phase tRF (19,350 peaks, FDR < 0.1; E, F) or with light-phase tRF (26,842 peaks, FDR < 0.1; G, H), each relative to *ad libitum* HFD. (I-L) Accessibility of representative genomic loci and motif-containing regions at ZT4 across HFD, HFDtRFDark, and HFDtRFLight conditions. Genome browser tracks at the *Dbp* locus (I, unit: CPM) and ATAC-seq signal at regions containing DBP (541 sites) and TEF (535 sites) motifs that lose accessibility under HFD compared to chow (J). Genome browser tracks at the *Smarca5* locus (K, unit: CPM) and ATAC-seq signal at regions containing ETS-family motifs that gain accessibility under HFD compared to chow (L). n=2 mice per time point per group. Color scales in (A, C, E, G) indicate z-scored chromatin accessibility across a 4-kb window centered on peak summits (−2 kb to +2 kb; 25-bp bins), normalized across all samples shown in each comparison. Color scales in (J, L) indicate z-scored chromatin accessibility at motif-containing regions, normalized across HFD, HFDtRFDark, and HFDtRFLight samples.
